# Cerebral cortical functional hyperconnectivity in a mouse model of spinocerebellar ataxia type 8 (SCA8)

**DOI:** 10.1101/2024.06.20.599947

**Authors:** Angela K. Nietz, Laurentiu S. Popa, Russell E. Carter, Morgan L Gerhart, Keerthi Manikonda, Laura P.W. Ranum, Timothy J. Ebner

## Abstract

Spinocerebellar Ataxia Type 8 (SCA8) is an inherited neurodegenerative disease caused by a bidirectionally expressed CTG●CAG expansion mutation in the *ATXN-8* and *ATXN8-OS* genes. While primarily a motor disorder, psychiatric and cognitive symptoms have been reported. It is difficult to elucidate how the disease alters brain function in areas with little or no degeneration producing both motor and cognitive symptoms. Using transparent polymer skulls and CNS-wide GCaMP6f expression, we studied neocortical networks throughout SCA8 progression using wide-field Ca^2+^ imaging in a transgenic mouse model of SCA8. We observed that neocortical networks in SCA8+ mice were hyperconnected globally which led to network configurations with increased global efficiency and centrality. At the regional level, significant network changes occurred in nearly all cortical regions, however mainly involved sensory and association cortices. Changes in functional connectivity in anterior motor regions worsened later in the disease. Near perfect decoding of animal genotype was obtained using a generalized linear model based on canonical correlation strengths between activity in cortical regions. The major contributors to decoding were concentrated in the somatosensory, higher visual and retrosplenial cortices and occasionally extended into the motor regions, demonstrating that the areas with the largest network changes are predictive of disease state.

## Introduction

Nucleotide repeat expansion mutations cause several neurological disorders, including Huntington’s Disease (HD), Myotonic Dystrophy (DM), and many of the spinocerebellar ataxias (SCAs; 1). Repeat expansion mutations are variable in size and for most disorders become both pathological and genetically unstable at a disease-specific repeat length threshold (2, 3). Spinocerebellar ataxia type 8 (SCA8) is an autosomal dominant inherited disease caused by a bidirectionally expressed nucleotide expansion mutation in the *ATXN8* and *ATXN8OS* genes. Onset of SCA8 in humans typically occurs in mid-adulthood and symptoms become progressively worse throughout the disease. There is a high degree of reduced penetrance with the SCA8 mutation, although most affected patients have repeat expansions > 70 CTG●CAGs (4). Phenotypically, SCA8 is characterized by unstable gait, dysarthria, nystagmus, and other motor symptoms (5). Clinical imaging reveals cerebellar atrophy in both the hemispheres and vermis with mild brain stem atrophy and neocortical atrophy in some cases (5–9).

At the molecular level, the CTG●CAG mutation is transcribed into CAG and CUG expansions in RNAs. These expansion RNAs undergo repeat-associated non-AUG (RAN) translation into polyserine, polyalanine, and polyglutamine expansion proteins (10, 11). Production of these homopolymeric RAN proteins can trigger apoptosis and lead to toxic gain of function effects (10, 12). The CUG expansion RNAs in SCA8 and DM can sequester muscleblind-like protein 1 (MBNL1), an RNA binding protein critical for brain structural integrity (13–15). In SCA8, CUG expansion RNAs cause increased expression of CUG triplet repeat RNA binding protein 1 - muscleblind-like protein 1 (CUGBP1-MBNL1) that regulates the CNS target, GABA-A transporter 4 (GAT4/GABT4). Increased levels of GABT4 are found in SCA8 mice and human patient tissue. These findings have a functional correlate in SCA8 mice of increased cerebellar neural responses to stimulation (2, 15), suggestive of decreased GABAergic inhibitory tone. Therefore, RAN protein intranuclear inclusions and RAN protein sequestration gain-of-function profoundly alter both brain structure and function.

As SCA8 is typically considered a cerebellar movement disorder, investigations into executive and cognitive function are limited. However, studies in which cognitive and psychiatric symptoms were self-reported, described SCA8 patients with personality changes, mood disturbances, anxiety, and depression (16). More recent work studying cognitive decline in SCA8 patients found attention and information processing deficits, including reduced detection of visual targets and reduced performance in the Stroop Color/Word Interference test (7). Additional deficits are found in executive function, and verbal tasks; however, memory was largely unaffected. Further, post-mortem brain tissue from SCA8 patients and a transgenic mouse model of SCA8 suggests brain-wide pathology. Cortical atrophy is seen in SCA8 patients on MRI and post-mortem protein abnormalities are found in the neocortex (9, 15). In both the mouse and human patients, RAN protein accumulation is not confined to the cerebellum and has been found in the motor cortex, brain stem, and white matter tracts (17). Therefore, both the pathological and clinical findings suggest that neocortical dysfunction in SCA8 is likely, as observed in other spinocerebellar ataxias (18–20). Also, these cortical changes could be manifest as changes in functional connectivity (FC), as observed in other neurological disorders (21–24). We would also expect extensive involvement of the motor cortices.

In addition to direct pathology, cortical dysfunction in SCA8 could also arise from disrupted input. The cerebellum has extensive reciprocal connections with the neocortex through the cerebello-thalamo-cortical and cortico-ponto-cerebellar pathways (25–27). This long-range cerebello-cortical loop could provide a neural substrate for propagation of cerebellar pathophysiology in SCA8 to neocortical networks. Here, we used neocortex-wide Ca^2+^ imaging to investigate changes in neocortical processing in a mouse model of SCA8. We find that the functional segmentation of the neocortex in SCA8+ mice is largely unchanged compared to non-transgenic controls (2). However, FC analysis revealed hyperconnectivity and stronger connections across atlas regions both before and after symptoms developed in SCA8 transgenic mice. These connectivity changes produced neocortical networks with more clustering, increased efficiency, and more network communities. SCA8 networks showed localized changes in the posterior sensory and sensory integration areas of the neocortex, in addition to the motor regions, which could be used to decode animal genotypes using a generalized linear model. These data suggest that neocortical processing is fundamentally altered prior to symptom onset in SCA8.

## Results

### Database and experimental paradigm

SCA8 transgenic mice (SCA8+; n = 7) and non-transgenic controls (NT control; n = 9) were imaged on a freely moving disc treadmill allowing for spontaneous rest and locomotion for ∼1 hour per session (∼10 trials; 5.5 mins per trial) throughout disease progression (see Methods; Figure 1A). Cortex-wide GCaMP6f expression was achieved using a retro-orbital injection and verified using post-hoc immunohistochemistry (Figure 1B). Neural activity was monitored in these mice throughout disease progression with disease onset defined as a 10% drop in pre-disease maximal weight (Figure 1C). Analysis was done using defined chronological epochs (see Methods; Figure 1C). Implanted transparent polymer skulls were aligned to the atlas using a multistep registration process. First Allen CCF landmarks were aligned to pre-craniotomy landmarks (Figure 1D). Next, the craniotomy path and implant border were aligned (Figure 1E). Finally, the Allen atlas was back transformed to the implanted window. The aligned implanted windows allowed visualization of layer II/III neocortical activity spanning the secondary motor cortex to the visual cortex in the anterior-posterior direction and the retrosplenial cortex to the medial edge of the barrel fields in the medial-lateral direction (Figure 1F).

**Figure 1:**
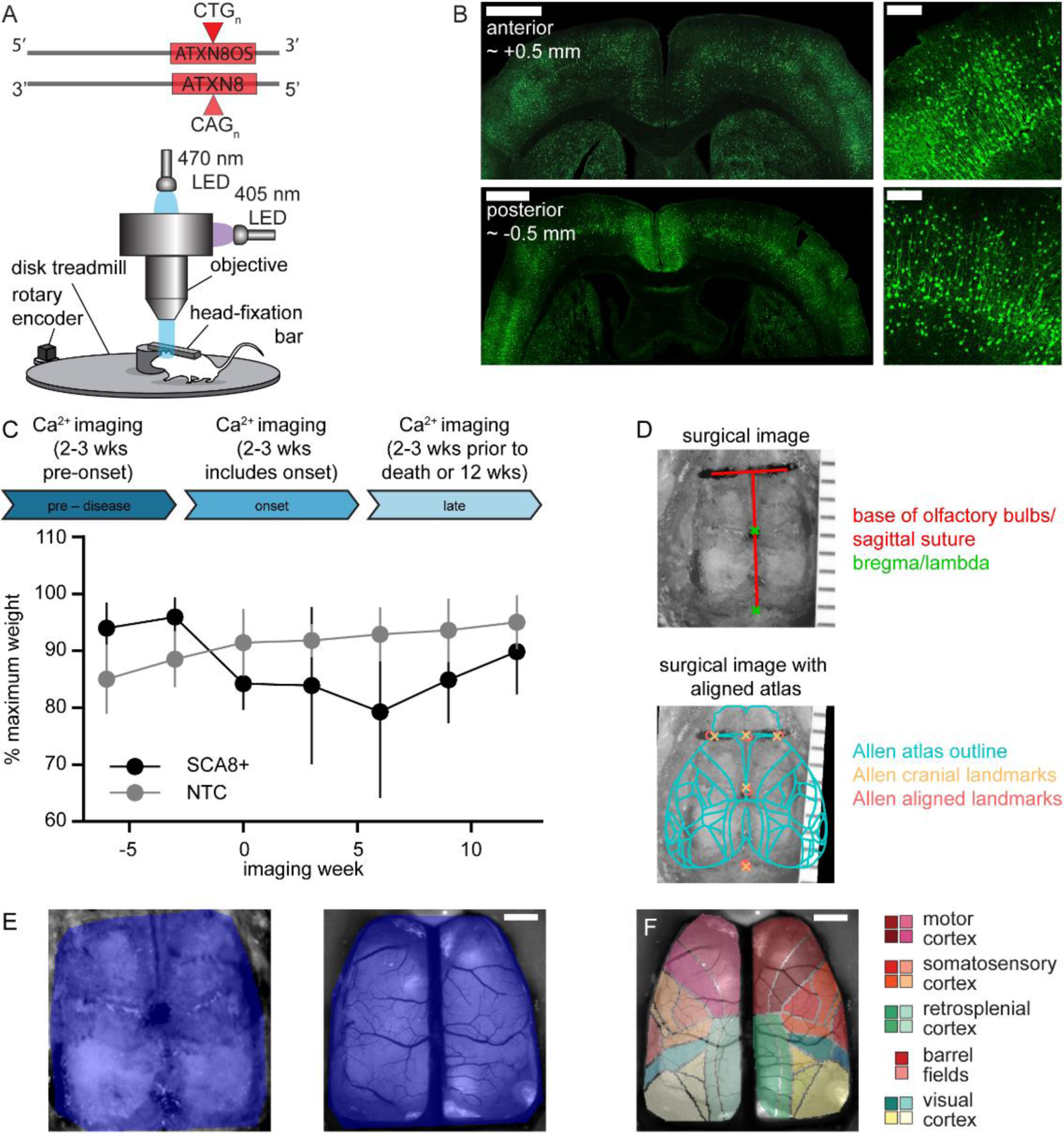
Viral GCaMP expression and wide-field Ca^2+^ imaging paradigm in SCA8+ mice. **A)** Schematic showing the expansion mutation in the human SCA8 transgene (top). Experimental setup showing Ca^2+^ imaging paradigm with animal head-fixed above a freely moving treadmill (bottom). **B)** Example coronal images of the neocortex showing broad expression of the Ca^2+^ sensor GCaMP6f after retro-orbital viral delivery (left; scale bars 1 mm). Example confocal image stacks showing GCaMP6f expression throughout the neocortical layers (right; scale bars 200 µm). **C)** Flow chart showing the imaging timeline across SCA8 disease progression and established analysis epochs (top) and a line graph showing animal weight (% maximum) across weeks of imaging used to empirically define SCA8 onset (black line - SCA8+; gray line - NT control; mean ± SD; week 0 - SCA8 onset). **D)** Example image of surgical craniotomy with anatomical landmarks (top; red - olfactory bulb base/inferior cerebral vein & sagittal suture; green - bregma & lambda; ruler ticks 1 mm). **D)** Image of surgical craniotomy aligned to the Allen CCF using defined anatomical landmarks (bottom; blue - Allen CCF outline; orange - Allen cranial landmarks; pink - aligned surgical cranial landmarks). **E)** Surgical (left) and neocortical imaging field of view (right) and their associated masks (dark blue) used for atlas alignment. **F)** Example image showing final alignment of Allen CCF to the Ca^2+^ imaging field of view (E-F scale bars 1 mm).

### SCA8 transgenic mice and controls show similar IC coverage across functional atlas areas

The first question addressed is whether functional segmentation of the cortex differed between SCA8+ and FVB NT control animals. Spatial independent component analysis (ICA) was run on a mouse level by concatenating data from the pre-disease, onset, and late phases (see Methods; 28) yielding a single set of independent components (ICs; Figure 2A). SCA8+ and NT control animals have similar numbers (Figure 2B; SCA8+: 55.3 ± 5.9; NT control: 50 ± 8.5; MW: p = 0.15, n = 7,9) and coverage of the ICs (SCA8+: 78.4 ± 3.7%; NT control: 74.2 ± 5.7%; MW: p = 0.17; n = 7,9). The ICs were assigned to CCF atlas regions based on their center positions (Figure 2C). Similar to the overall IC values between SCA8+ and NT controls, each atlas region contains similar numbers of ICs (Figure 2D; for statistical details see Table 1). The numbers and spatial distribution of ICs imply that the functional segmentation remains intact in SCA8 mice. For each IC, the hemodynamic-corrected ΔF/F time series was extracted for both SCA8+ and NT controls at each phase of disease progression. All phases of disease in both genotypes show GCaMP fluorescence transients of varying amplitudes, indicative of neuronal activity; similar to previous reports (29–31). Example time series from select ICs show the expected fluorescence modulation with comparable levels (Figure 2E). Qualitatively, the fluorescence signals suggest stronger correlations between cross regional ICs in the SCA8+ mice compared to the NT controls. The subsequent functional connectivity (FC) analyses address how these patterns of correlation across the neocortex are organized and how they differ between genotypes.

**Figure 2:**
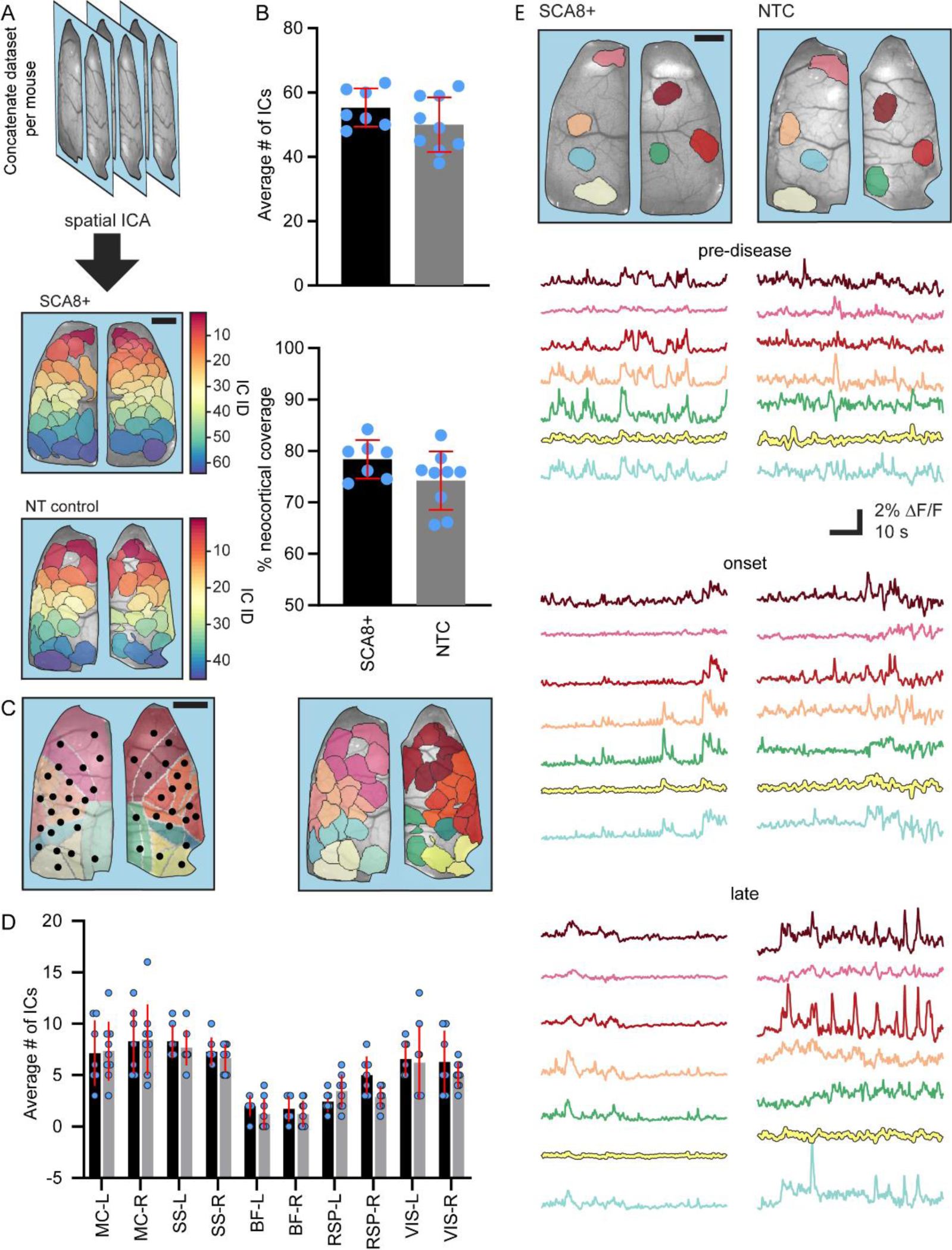
SCA8+ and NT control mice show similar neocortical functional segmentations using spatial ICA. **A)** Schematic showing chronological concatenation of data over 3 defined SCA8 disease phases (top) and the resulting neocortical functional segmentations for an SCA8+ (middle) and NT control mouse (bottom; different colors denote individual ICs). **B)** Scatter bar plots showing the average number of ICs (top) per SCA8+ (black) or NT control (gray) mouse after spatial ICA processing and average percentage of cortical coverage by ICs (bottom). **C)** Example neocortical images showing aligned pseudo-colored Allen CCF overlay and IC centroids (left) used to assign and color code ICs to functional atlas regions (right). Scale bars 1 mm. **D)** Scatter bar plot showing the average number of ICs assigned to each functional atlas region for SCA8+ (black) and NT control (gray) animals. MC - Motor Cortex; SS - Somatosensory Cortex; BF - Barrel Field Cortex; RSP- Retrosplenial Cortex; VIS - Visual Cortex; R - right; L - left. **E)** Images showing select ICs (colored areas) overlayed on the neocortex for an SCA8+ (left) and NT control (right) mouse and examples of their corresponding Ca^2+^ traces for each of the three disease phases (bottom).

**Table 1.** Numerical comparison of ICs per CCF functional area.

| **Atlas Area** | **# ICs per region**<br>**(mean ± SD)** | **MW p-value**<br>**(SCA8+ vs.**<br>**NTC)** |
| --- | --- | --- |
| **MC_L** | SCA8+: 7.14 ± 3.18<br>NTC: 7.33 ± 2.87 | p > 0.99 |
| **MC_R** | SCA8+: 8.29 ± 3.15<br>NTC: 8.44 ± 3.43 | p > 0.99 |
| **SS_L** | SCA8+: 8.29 ± 1.6<br>NTC: 7.67 ± 1.73 | p > 0.99 |
| **SS_R** | SCA8+: 7.29 ± 1.38<br>NTC: 6.67 ± 1.32 | p > 0.99 |
| **BF_L** | SCA8+: 2.00 ± 1.00<br>NTC: 1.22 ± 1.48 | p = 0.90 |
| **BF_R** | SCA8+: 1.71 ± 1.25<br>NTC: 1.22 ± 1.30 | p > 0.99 |
| **RSP_L** | SCA8+: 2.43 ± 0.98<br>NTC: 3.44 ± 1.59 | p = 0.90 |
| **RSP_R** | SCA8+: 5.00 ± 1.83<br>NTC: 2.89 ± 1.05 | p = 0.21 |
| **VIS_L** | SCA8+: 6.57 ± 1.72<br>NTC: 6.22 ± 3.60 | p > 0.99 |
| **VIS_R** | SCA8+: 6.29 ± 3.04<br>NTC: 5.00 ± 1.22 | p > 0.99 |

Average number of ICs ( ± SD) and statistical comparisons per CCF atlas region between SCA8+ and NT control mice. SCA8 - SCA8+; NTC - NT control; MW - Mann-Whitney test; MC - motor cortex; SS - somatosensory; BF - barrel fields; RSP - retrosplenial cortex; VIS - visual cortex; L - left; R - right.

### Global network functional connectivity is disrupted in SCA8+ mice

For each disease phase, FC was assessed by correlating IC average fluorescence signals within each atlas region to other atlas regions using canonical correlation analysis (CCA; 32, 33). The CCA matrices were thresholded and all values <0.5 were set to zero and graphed using Matlab (*graph*, *plot*). The FC graphs were plotted over brain images with the nodes at the center of each atlas area and node size signifying relative connection density. The edges are CCA values ≥0.5 with edge color signifying the weight for each connection. Examples of atlas assigned ICs from individual SCA8+ animals (Figure 3A-B) and NT control animals (Figure 3D-E) show the larger atlas cortical areas (e.g., motor, somatosensory, visual, retrosplenial) are represented in both genotypes. Additionally, the size, number, and spatial distribution of ICs identified in both genotypes show that functional parcellation of the neocortex is much more fine-grained than suggested by the CCF. Taken together, these data show that the structure and organization of the functional parcellation is not fundamentally altered in SCA8+ mice.

**Figure 3:**
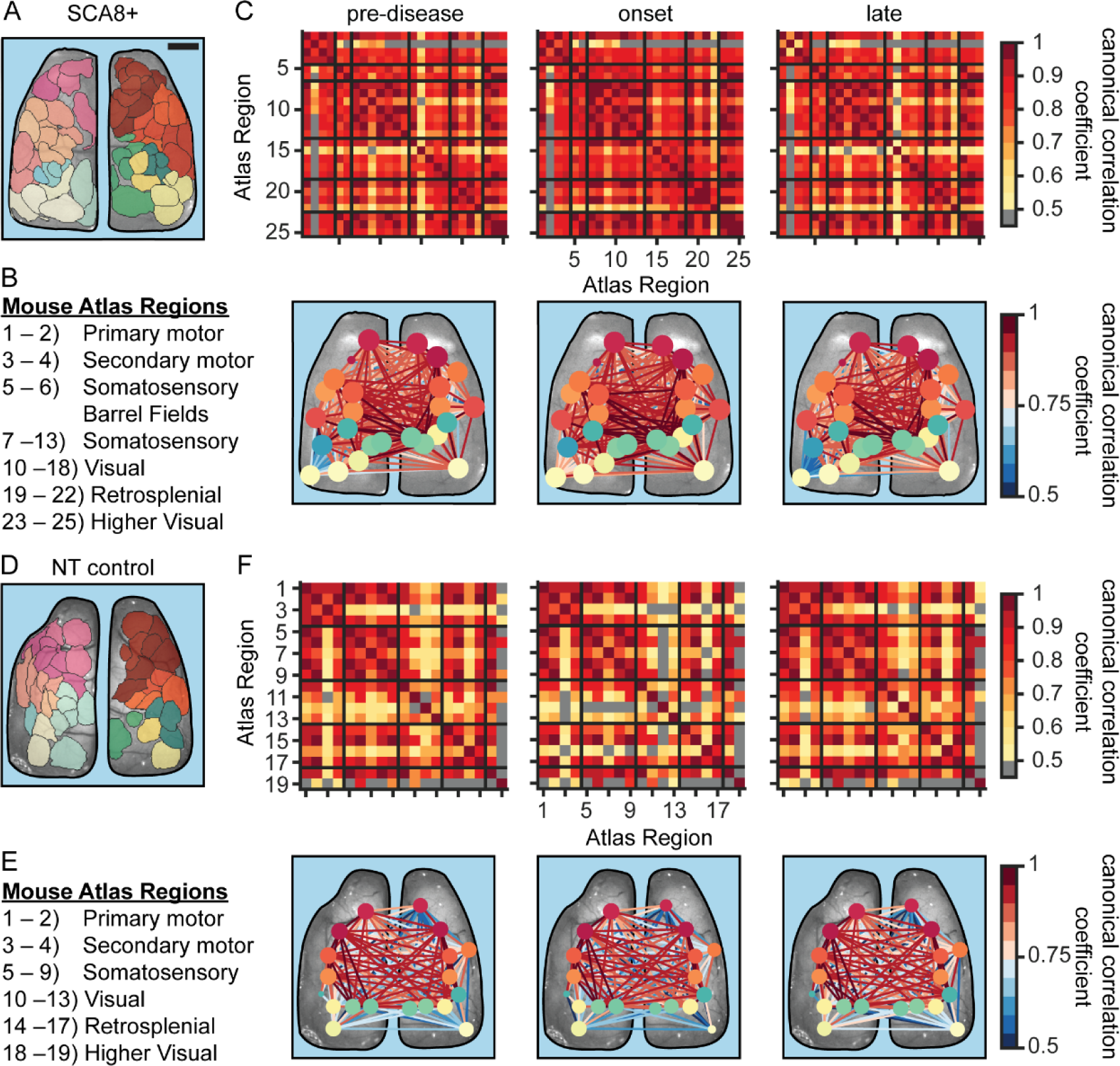
SCA8+ mice have hyperconnected neocortical networks. **A,D)** Example ICs from an SCA8+ (A) or NT control (D) mouse color-coded to their assigned atlas regions. **B,E)** Numerical IDs for broad Allen CCF atlas regions corresponding to the matrices shown in C and F, respectively. **C,F)** Average thresholded (≥0.5) canonical correlation analysis (CCA) matrices (top) for the atlas regions defined in B or E for each disease phase in an SCA8+ (C) and NT control (E) mouse. Canonical correlations are calculated using the Ca^2+^ signals of the ICs belonging to each atlas region keeping only the first canonical correlate for each comparison. Black lines delineate major CCF functional divisions. Corresponding neocortical network graphs (bottom) for the CCA matrices shown above. Nodes and node colors represent CCF atlas areas and edges represent functional connections assessed by CCA. Node size indicates relative number of connections and edge color indicates relative connection strength.

Individual SCA8+ animals show a global increase in the canonical correlation between Ca^2+^ signals in the pre-disease phase that persists throughout disease progression (Figure 3C, top) compared to NT controls (Figure 3F, top). These global increases in correlation produce an increased number and strength of connections in network graphs, altering the terrain of the FC maps, with the increased connectivity observed across SCA8+ animals compared to NT controls (Figure 4A). The canonical correlations of IC fluorescence signals in SCA8+ animals are greater both within and between atlas regions (Figure 4B, top) compared to NT controls (Figure 4C, bottom) and the increase is widespread. Therefore, the SCA8+ mice have greater intra- and inter-nodal FC than NT controls.

**Figure 4:**
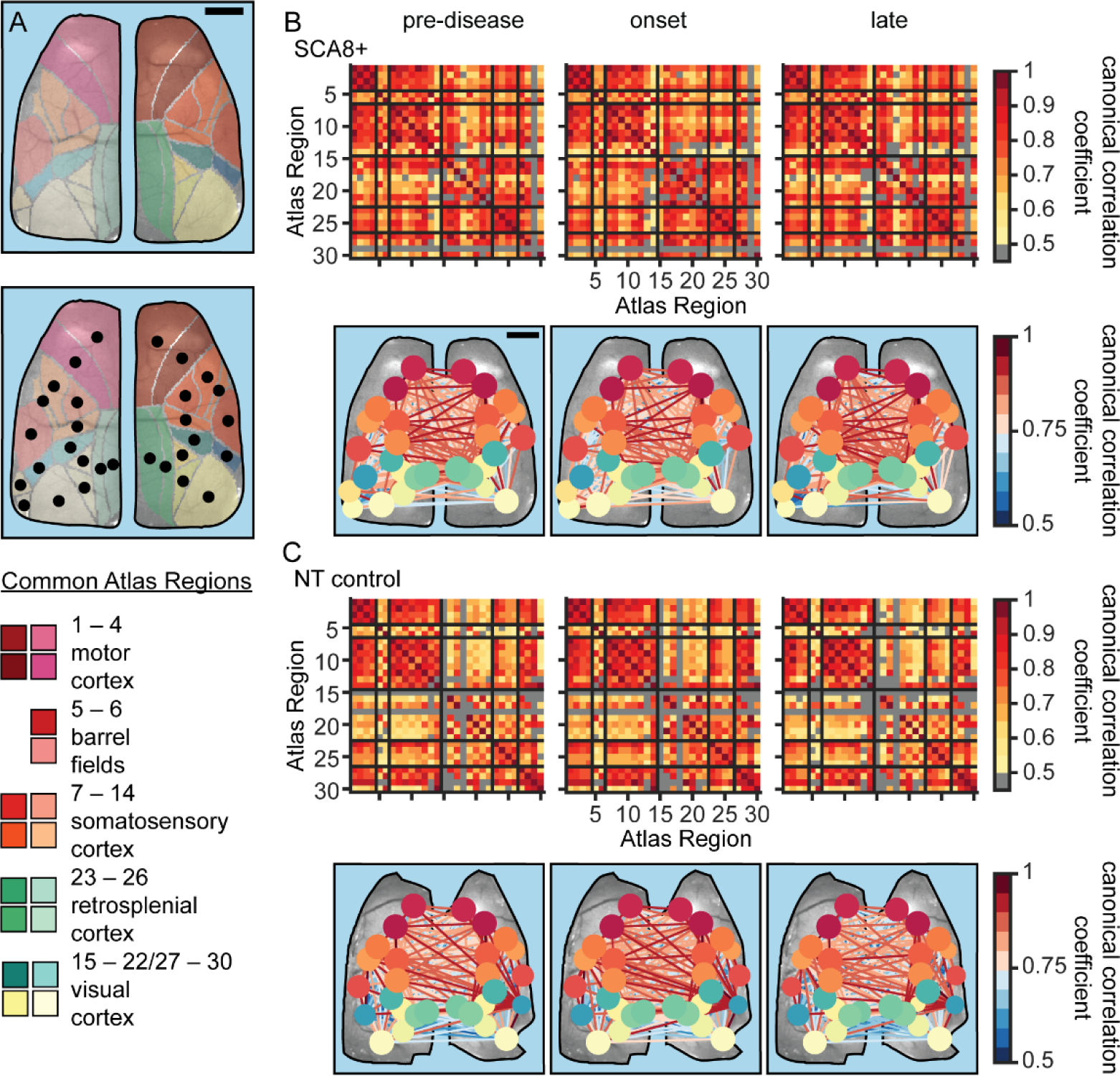
Neocortical networks are globally hyperconnected across SCA8+ mice. **A)** Pseudo-colored Allen CCF showing atlas regions visible across all subjects (top, 30 total regions) and the centroid of each atlas region (middle; centroids denoted by black dots). Bottom panel shows color-coded legend for visible atlas regions above and their numerical IDs corresponding to the matrices shown in B and C. **B-C)** Average CCA matrices (top) across all SCA8+ (B) and all NT control (C) mice for all visible atlas regions in A at each disease phase (gray squares are connections below the CCA threshold of 0.5) and their associated network graphs (bottom). Black lines denote major CCF functional divisions. Corresponding neocortical network graphs (bottom) for the CCA matrices shown above. Nodes and node colors represent CCF atlas areas and edges represent functional connections assessed by CCA. Node size indicates relative number of connections and edge color indicates relative connection strength. Missing nodes denote regions that had no correlations above the set threshold.

We quantified these network-wide changes in FC using the Brain Connectivity Toolbox (34, 35). As observed in the canonical correlation matrices (Figures 3 and 4), the average connection density and global nodal strength are significantly higher in the SCA8+ animals compared to NT controls during all three disease phases (Figure 5A left, middle; for statistical details see Table 2), suggesting a hyperconnectivity and strongly coupled cross-regional fluorescence signals. In all three disease phases, the average global efficiency, global eigenvector centrality, and number of community partitions are also increased in SCA8+ animals (Figure 5A right; Figure 5B left, middle; also see Table 2) compared to NT controls. Global efficiency measures the ease of information exchange across the network and eigenvector centrality measures the magnitude of influence a node has over network processing. Interestingly, the average global transitivity (an analog of the clustering coefficient which measures connectivity of a node to its neighbors) is significantly increased only during the early and late disease phases (Figure 5B, right; also see Table 2). As several global connectivity and network topology measures are altered in SCA8+ animals compared to NT controls prior to our empirical definition of SCA8 onset, this suggests that neocortical alterations happen early-on in disease before the appearance of overt physical symptoms.

**Figure 5:**
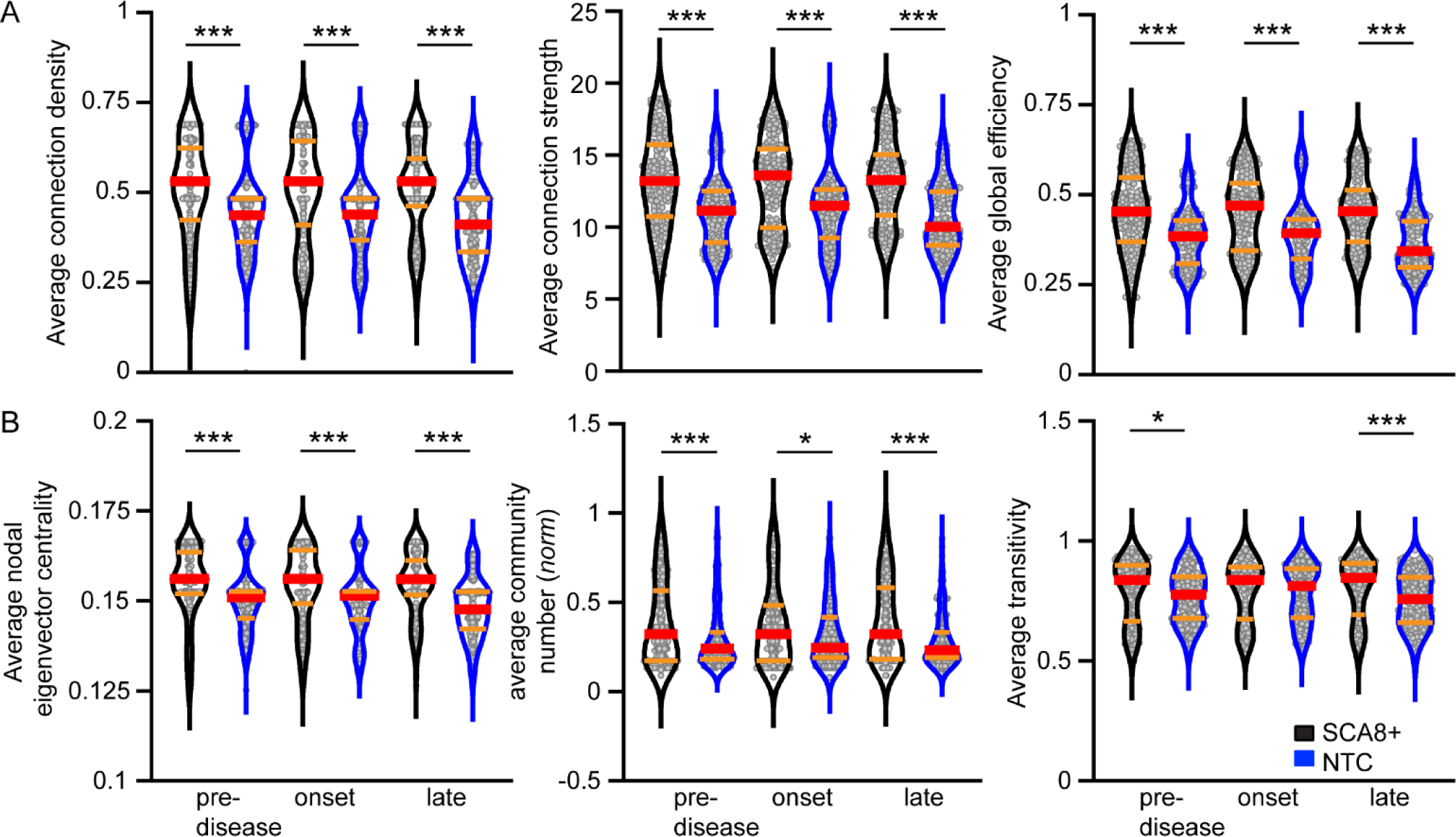
Network configuration is altered across disease phases in SCA8+ mice. **A)** Violin plots showing the average network density (left), average strength per node (middle), and global network efficiency (right) per trial in SCA8+ (black) and NT control (blue) animals at each disease phase. **B)** Violin plots showing average eigenvector centrality per node (left), average number of communities (middle), and average network transitivity (right) per trial in SCA8+ and NT control animals. Red bars denote median and orange bars denote the quartiles. Community partitions were calculated using only regions present in each mouse and normalized to the number of nodes in each animal’s network to allow equal comparison. Black bars denote statistically significant differences between SCA8+ and NT control mice according to a 2-way ANOVA with Bonferroni post-hoc comparisons matching the statistics presented in Table 2. *** p < 0.0001; * p < 0.05.

**Table 2.**
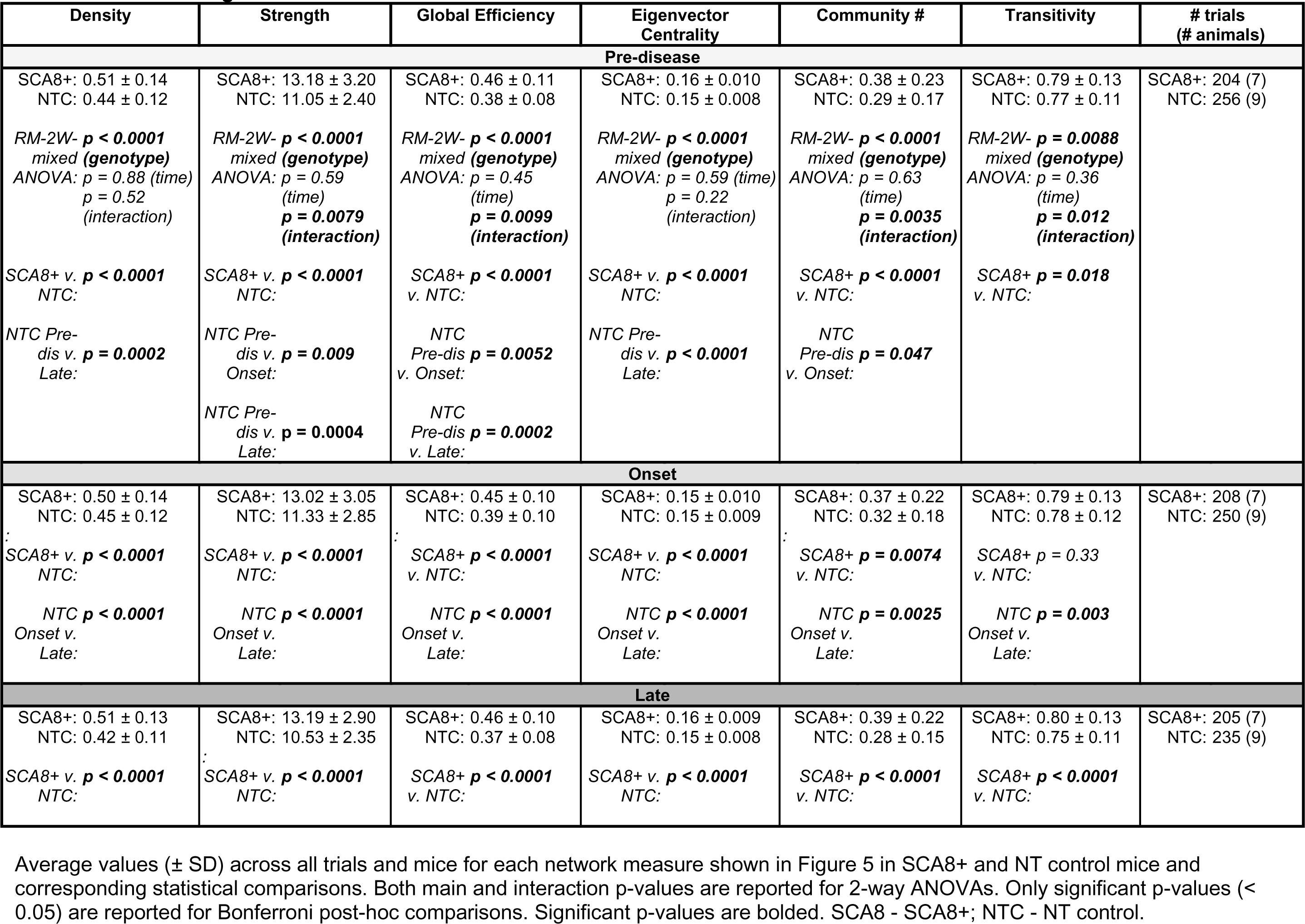
Network configuration measures in SCA8 and NT control mice.

| Density | Strength | Global Efficiency | Eigenvector Centrality | Community # | Transitivity | # trials (# animals) |
| --- | --- | --- | --- | --- | --- | --- |
| **Pre-disease** |  |  |  |  |  |  |
| SCA8+: 0.51 ± 0.14<br>NTC: 0.44 ± 0.12<br>*RM-2W- **p < 0.0001***<br>*mixed (genotype)*<br>*ANOVA: p = 0.88 (time)*<br>*p = 0.52 (interaction)* | SCA8+: 13.18 ± 3.20<br>NTC: 11.05 ± 2.40<br>*RM-2W- **p < 0.0001***<br>*mixed (genotype)*<br>*ANOVA: p = 0.59 (time)*<br>***p = 0.0079***<br>*(interaction)* | SCA8+: 0.46 ± 0.11<br>NTC: 0.38 ± 0.08<br>*RM-2W- **p < 0.0001***<br>*mixed (genotype)*<br>*ANOVA: p = 0.45 (time)*<br>***p = 0.0099***<br>*(interaction)* | SCA8+: 0.16 ± 0.010<br>NTC: 0.15 ± 0.008<br>*RM-2W- **p < 0.0001***<br>*mixed (genotype)*<br>*ANOVA: p = 0.59 (time)*<br>*p = 0.22 (interaction)* | SCA8+: 0.38 ± 0.23<br>NTC: 0.29 ± 0.17<br>*RM-2W- **p < 0.0001***<br>*mixed (genotype)*<br>*ANOVA: p = 0.63 (time)*<br>***p = 0.0035***<br>*(interaction)* | SCA8+: 0.79 ± 0.13<br>NTC: 0.77 ± 0.11<br>*RM-2W- **p = 0.0088***<br>*mixed (genotype)*<br>*ANOVA: p = 0.36 (time)*<br>***p = 0.012***<br>*(interaction)* | SCA8+: 204 (7)<br>NTC: 256 (9) |
| SCA8+ v. NTC: **p < 0.0001**<br>NTC Pre-dis v. Late: **p = 0.0002** | SCA8+ v. NTC: **p < 0.0001**<br>NTC Pre-dis v. Onset: **p = 0.009**<br>NTC Pre-dis v. Late: **p = 0.0004** | SCA8+ v. NTC: **p < 0.0001**<br>NTC Pre-dis v. Onset: **p = 0.0052**<br>NTC Pre-dis v. Late: **p = 0.0002** | SCA8+ v. NTC: **p < 0.0001**<br>NTC Pre-dis v. Late: **p < 0.0001** | SCA8+ v. NTC: **p < 0.0001**<br>NTC Pre-dis v. Onset: **p = 0.047** | SCA8+ v. NTC: **p = 0.018** |  |
| **Onset** |  |  |  |  |  |  |
| SCA8+: 0.50 ± 0.14<br>NTC: 0.45 ± 0.12<br>SCA8+ v. NTC: **p < 0.0001**<br>NTC Onset v. Late: **p < 0.0001** | SCA8+: 13.02 ± 3.05<br>NTC: 11.33 ± 2.85<br>SCA8+ v. NTC: **p < 0.0001**<br>NTC Onset v. Late: **p < 0.0001** | SCA8+: 0.45 ± 0.10<br>NTC: 0.39 ± 0.10<br>SCA8+ v. NTC: **p < 0.0001**<br>NTC Onset v. Late: **p < 0.0001** | SCA8+: 0.15 ± 0.010<br>NTC: 0.15 ± 0.009<br>SCA8+ v. NTC: **p < 0.0001**<br>NTC Onset v. Late: **p < 0.0001** | SCA8+: 0.37 ± 0.22<br>NTC: 0.32 ± 0.18<br>SCA8+ v. NTC: **p = 0.0074**<br>NTC Onset v. Late: **p = 0.0025** | SCA8+: 0.79 ± 0.13<br>NTC: 0.78 ± 0.12<br>SCA8+ v. NTC: **p = 0.33**<br>NTC Onset v. Late: **p = 0.003** | SCA8+: 208 (7)<br>NTC: 250 (9) |
| **Late** |  |  |  |  |  |  |
| SCA8+: 0.51 ± 0.13<br>NTC: 0.42 ± 0.11<br>SCA8+ v. NTC: **p < 0.0001** | SCA8+: 13.19 ± 2.90<br>NTC: 10.53 ± 2.35<br>SCA8+ v. NTC: **p < 0.0001** | SCA8+: 0.46 ± 0.10<br>NTC: 0.37 ± 0.08<br>SCA8+ v. NTC: **p < 0.0001** | SCA8+: 0.16 ± 0.009<br>NTC: 0.15 ± 0.008<br>SCA8+ v. NTC: **p < 0.0001** | SCA8+: 0.39 ± 0.22<br>NTC: 0.28 ± 0.15<br>SCA8+ v. NTC: **p < 0.0001** | SCA8+: 0.80 ± 0.13<br>NTC: 0.75 ± 0.11<br>SCA8+ v. NTC: **p < 0.0001** | SCA8+: 205 (7)<br>NTC: 235 (9) |

### Area specific alterations in functional connectivity in SCA8

We hypothesized that alterations in SCA8 networks would be concentrated in the motor cortices as SCA8 is primarily considered a motor disorder. To determine if there are location-specific alterations in the SCA8+ networks, node-specific analyses were performed using the major anatomical regions and all sub-parcellations within major regions were considered significantly different if the Tukey-adjusted p-value was < 0.05. Contrary to our working hypothesis, many changes occur in non-motor areas in the posterior cortex, including the visual and retrosplenial areas, in addition to the sensorimotor cortices. Nodal degree is increased in the secondary motor cortices, retrosplenial cortices, and subregions of the visual cortices (Figure 6A; 3-way ANOVA p < 0.0001 *genotype/time/atlas area/genotype & atlas area/genotype & time;* p = 0.002 *time & atlas area*; p = 0.033 *genotype & time & atlas area*). These increases in nodal degree persist from the pre-disease phase to the late disease phase and progress into the somatosensory and higher visual cortices in late disease. Most areas of the neocortex showed stable increases in connection strength with other cortical regions (Figure 6B; 3-way ANOVA p < 0.0001 *genotype/atlas area/time/genotype & time/genotype & atlas area*). These alterations persist in all disease phases demonstrating that network changes are most prominent in the sensory and association cortices.

**Figure 6:**
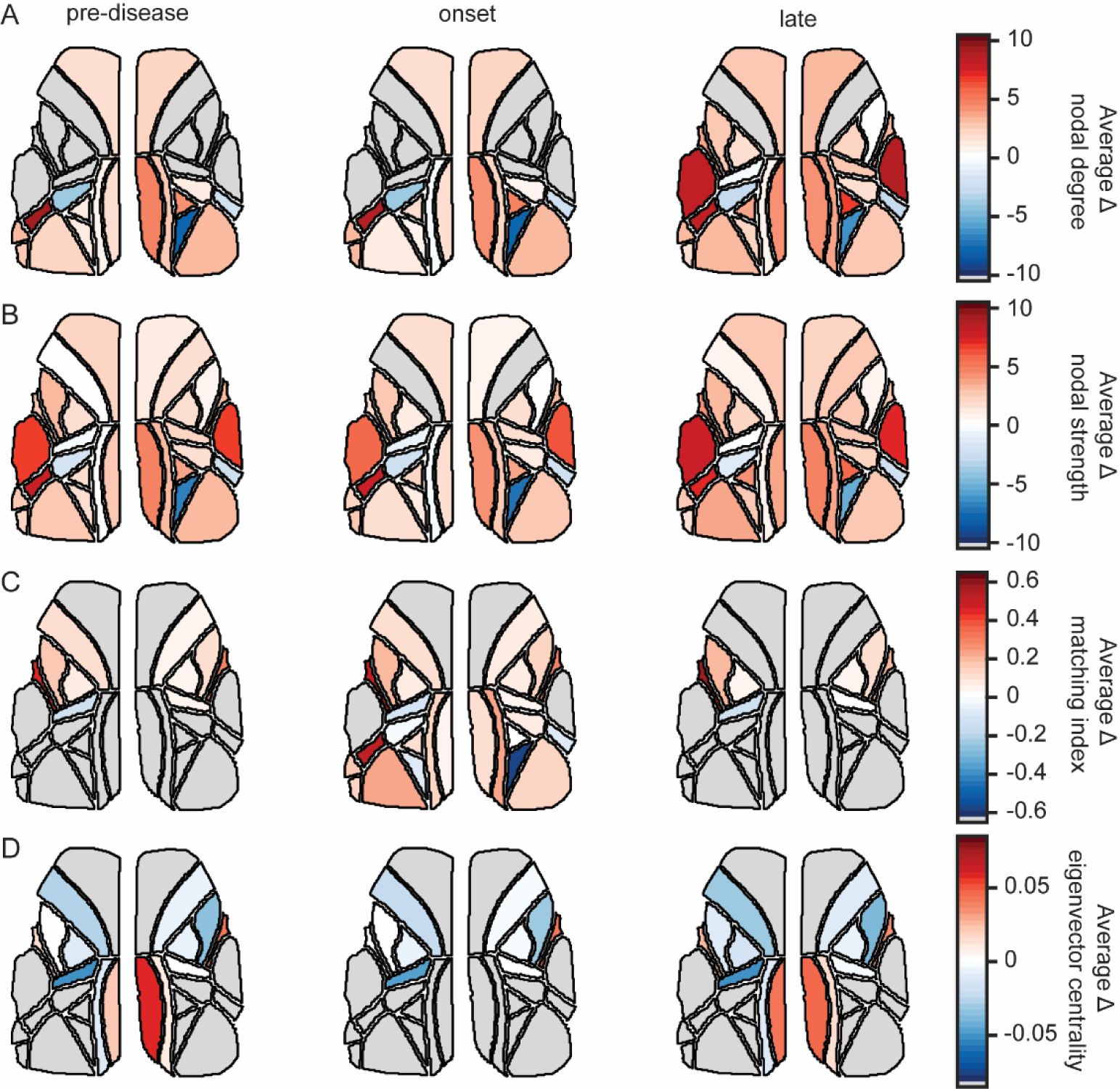
Region-specific changes in network topology drive global differences in SCA8 networks. Pseudo-colored Allen CCFs showing statistically significant changes in nodal degree **(A)**, strength **(B)**, matching index **(C)**, and eigenvector centrality **(D)** between SCA8+ mice and NT control mice for each disease phase. The magnitude of change in network measures for pseudo-coloring and scaling is calculated by subtracting the average measure across NT control animals for each atlas region from the average across SCA8+ animals. Positive changes in magnitude (red) indicate the network measure is increased SCA8+ animals compared to NT controls. Negative changes in magnitude (blue) indicate the network measure is reduced in SCA8+ mice compared to NT controls. All colored regions were significantly different between genotypes for the major atlas area they belong to (p < 0.05) with Tukey HSD post-hoc comparisons following a 3-way repeated measures mixed model ANOVA. All sub-regions within major atlas areas were considered significant if the Tukey test for the comparison was significant. All gray regions did not show significant differences in network measures per major atlas regions between genotypes.

Next, we examined whether specific nodes in the SCA8+ networks were more functionally similar to one another compared to NT control networks. To do this, we calculated the matching index for each node which measures connectivity overlap between two nodes (or redundancy of connections) which are not connected to one another (36). We found the networks in SCA8+ animals have an increased matching index compared to NT controls in the limb and trunk sensory cortices across disease phases (Figure 6C; 3-way ANOVA p < 0.0001 *atlas area/genotype & atlas area/genotype & time*; p = 0.0037 *genotype*). Matching index is selectively increased in the retrosplenial and visual areas of SCA8+ mice in the onset phase. These data reinforce the findings that specific sensory modalities and integrational areas contain many of the changes in FC in SCA8+ compared to NT control animals. Finally, we assessed nodal changes in eigenvector centrality in SCA8+ and NT control networks (Figure 6D; 3-way ANOVA p < 0.0001 *genotype/atlas area/genotype & atlas area/time & atlas area*; p =0.022 *time*; p = 0.00062 *genotype & time & atlas area*). Centrality is selectively reduced in the primary motor cortex and major somatosensory areas. In contrast, centrality is increased in the retrosplenial areas during the pre-disease and late phases. While this contrasts with the increase in global network centrality in SCA8+ animals, we hypothesize that the increase in global network centrality is driven by low magnitude increases in centrality across nodes which, when examined at an individual level, do not result in region specific significant centrality changes. These node specific alterations suggest that cortical areas with high sensory integration have considerable control over network processing in these SCA8 mice, more so than sensorimotor areas.

### Predicting SCA8 genotype using functional connectivity

Finally, we utilized CCA-based FC matrices to determine whether FC network properties can be used to accurately decode the genotype of animals using a stepwise generalized linear model (GLM). In our GLM, Bayesian information criterion (BIC) was used to determine the best-fit model and the minimum number of functional connections needed to predict animal genotype. At each phase of disease, the GLM decodes genotype with >98% accuracy, >98% precision, and >98.5% recall (Figure 7A; n = 10 cross-validations). When the model was trained using shuffled genotypes, model accuracy dropped significantly with < 53% accuracy, <46% precision, and <22% recall (Figure 7B; total n = 5000; iterations - 1000; cross-validations/shuffle - 5). These data show that FC can be used as a distinguishing feature for SCA8 disease prior to symptom onset in our model.

**Figure 7:**
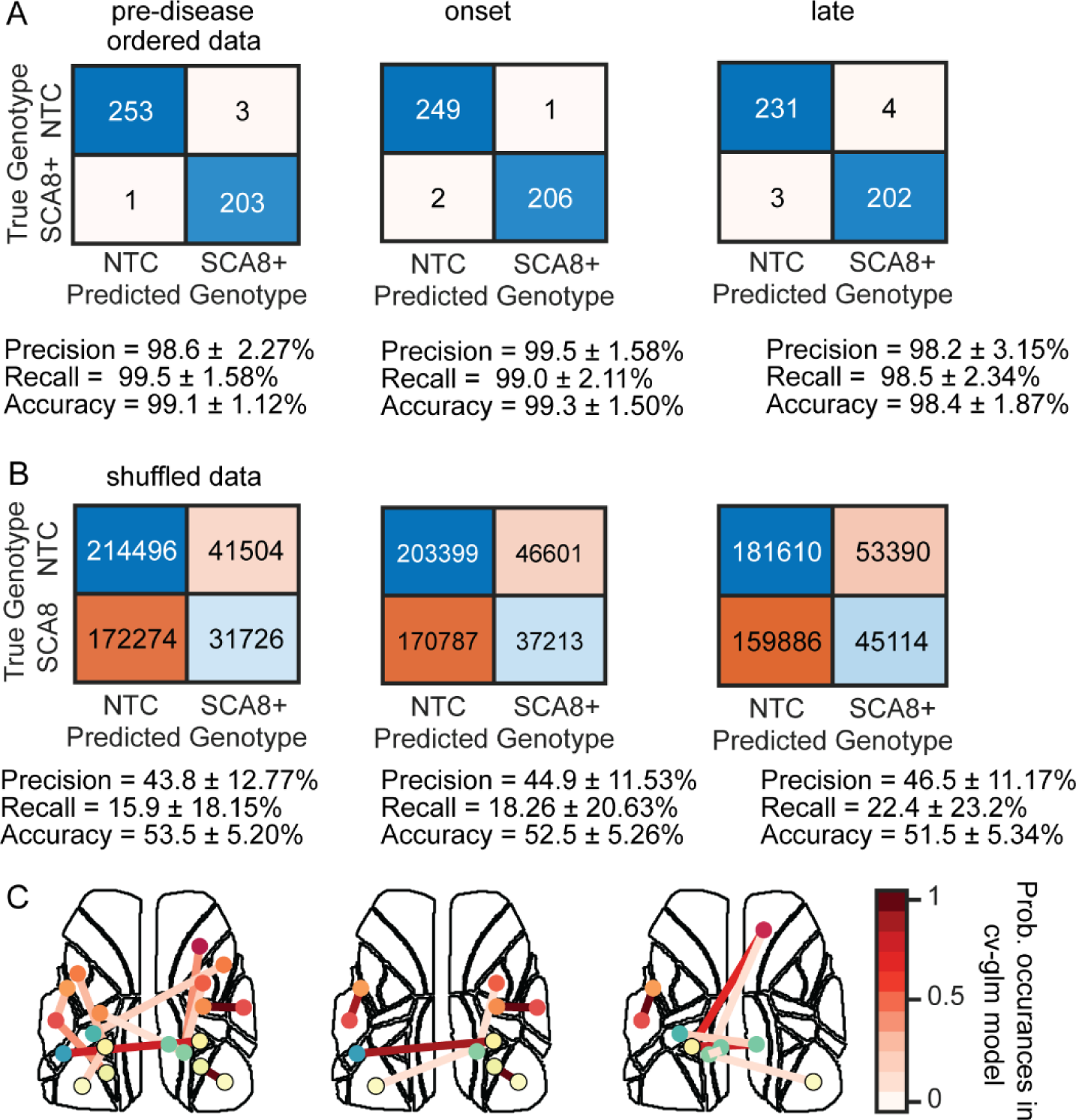
Functional connectivity can be used to decode genotype in SCA8 animals. **A)** Summative confusion charts (top) for each disease phase showing mouse genotype classification after training a stepwise GLM with ten-fold cross-validation and the average performance metrics associated with each GLM (mean ⨦ SD). **B)** Summative confusion charts (top) for disease phase showing mouse genotype classification after training a stepwise GLM with shuffled classification labels (1000 permutations each with 5-fold cross-validation) and their corresponding performance metrics (bottom). **C)** Allen CCF outlines and network nodes (color-coded to CCF atlas region) showing connections important for discriminating between genotypes in each GLM at each disease phase. Edges show important connections and color shows each connection’s probability of use across the 10 cross-validations of the GLM. Note that most of these areas had significant network measure changes in Figure 6.

Next, the GLM was used to determine which predictors (FC between atlas areas) were most effective at discriminating between SCA8+ mice and NT controls. The predictors repeatedly selected by the stepwise GLM algorithm across cross-validations provided the most predictive power. The most powerful predictive interactions are located primarily in the posterior sensory cortices and integration areas across all disease phases, particularly between the barrel and somatosensory cortices as well as the visual and retrosplenial cortices (Figure 7C). In the pre-disease and late disease stages, interactions between the motor cortices and retrosplenial areas contribute to the prediction of disease state. Interestingly, the areas most effective at predicting animal genotype are the same areas with specific changes in network connectivity metrics. These data show that neocortical network topology and sensorimotor integrational FC is fundamentally impaired throughout the lifespan of SCA8+ mice, and the altered topology provides highly robust information on the genotype.

## Discussion

We performed longitudinal neocortex-wide Ca^2+^ imaging in mice expressing the human SCA8 expansion mutation as well as non-transgenic control animals. To assess the FC between cortical areas, we used spatial ICA to functionally segment the cortex in an unbiased, data-driven manner and mapped the ICs onto the CCF. This was followed by CCA between atlas regions as the basis for determining differences in cortical network FC. Compared to the control animals, SCA8+ mice exhibit several global differences, including hyperconnectivity and increased connection magnitude as well as increases in efficiency, global eigenvector centrality, and number of communities. Interestingly, these changes are evident prior to our definition of the disease onset, a 10% loss of pre-onset maximal body weight. These global changes suggest that in the SCA8+ neocortex information processing is more fragmented and the specificity of information transfer across the network is impaired. At the regional (nodal) level, FC changes were evident not only in the motor cortices but also the posterior sensory and sensory integration regions. The most prominent changes over time are connection density and strength, with subtler changes in nodal matching index and nodal eigenvector centrality. Additionally, the region-specific changes increased, both spatially and in magnitude, including more sensory and higher visual regions as the disease progressed. The information in these region-specific changes could be used to perform near perfect decoding of animal genotypes using a generalized linear model.

### Neocortical functional segmentation is preserved in SCA8

Many studies of FC utilize canonical neocortical segmentations such as the Allen CCF or pool segmentations across animals (37, 38). This includes using the same segmentation for both control and models of disease (22, 38, 39). While common segmentations make interpretation easier, inter-subject variability and disease-related changes are not available. As neurodegenerative diseases can dramatically alter both brain structure and function, including in SCA8 (2, 17, 40), we could not presume that neocortical functional segmentation remains stable. Therefore, we used spatial ICA to determine if abnormalities exist in the functional segmentation of SCA8+ mice. We found that neocortex functional segmentation does not differ between SCA8+ and NT controls, with similar numbers and spatial distributions of ICs within the canonical CCF. The present finding of a stable functional segmentation is consistent with our previous results using a mouse model of mild traumatic brain injury (31) and that activity-dependent segmentations from spatial ICA are largely stable within subjects across time and behavior (28). These data suggest that neocortical functional segmentation remains stable even in this mouse model of cerebellar dysfunction.

### Neocortical networks are hyperconnected in SCA8+ mice

Using the Brain Connectivity Toolbox (34), we constructed FC networks and found cortical networks are globally hyperconnected in SCA8+ animals compared to NT controls, including increased connection density and strength. The network hyperconnectivity preceded our definition of disease onset and presentation of phenotypic symptoms. As the SCA8 mutation is constitutively expressed, pre-symptomatic effects on brain function are not necessarily unexpected. As disease progressed and motor symptoms became severe, global FC in the neocortex did not change appreciably between SCA8+ and NT controls. These findings suggest that FC alterations are not simply due to increasing behavioral deficits or health decline, but instead to early and persistent changes in SCA8+ brain function.

While we are not aware of comparable FC studies in SCA8 mouse models or patients, pre-symptomatic altered FC occurs in other neurodegenerative diseases. For example, several AD mouse models exhibit hyperexcitability in cortical and hippocampal networks and increased seizure susceptibility prior to the manifestation of memory deficits (41, 42). Frontal-cerebellar FC is reduced in Fredrich’s Ataxia patients whereas FC between cortical regions is enhanced (43). In SCA3 patients, cerebello-cerebral FC is disrupted and correlates with trinucleotide repeat length (44). Here, we show for the first time functional hyperconnectivity in a mouse model of SCA8.

### Cortical network topology is altered in SCA8

Investigation into network structure in SCA8+ and NT controls revealed different network configurations. Network efficiency and nodal centrality were increased in SCA8+ mice relative to NT controls and the network was partitioned into more, smaller communities with more clustered connectivity. Other disease states, like temporal lobe epilepsy, show similarly altered cortical networks (23, 24). Epileptic children display hyper-clustered, highly efficient network configurations, whereas adults display hyper-clustered but less efficient networks compared to non-epileptic individuals. In human stroke patients and two mouse stroke models, networks show hyperconnectivity, increased nodal strength, increased clustering of nodes in the network, and shorter characteristic path length (indicative of increased efficiency; 45). Individuals with Aꞵ deposits, suggestive of early AD, show brain area dependent increases or decreases in eigenvector centrality (46). In Huntington’s Disease (HD), resting state networks shift toward within-network hyperconnectivity but reduced connectivity between networks (47), and area-specific changes in FC are apparent with concurrent changes in network configuration (21). Finally, a zebrafish model of depression shows increased network modularity with more anatomically distributed communities (48), suggesting both a less structured organization and reduced long-range information transfer (49). In our SCA8+ mice, we propose that the increased network efficiency and global centrality together with the larger community number and clustering produce a more randomly distributed network where information flow is less segregated and causes disintegration of network processing.

### Region-specific hubs drive global network alterations in SCA8

Next, we evaluated whether the FC changes were uniform or area specific. Surprisingly, we found that in the pre-disease and onset disease phases, nodal degree and strength were increased in the secondary motor, visual, and retrosplenial cortices. In late disease, changes progressed to include the barrel, primary, motor, and major somatosensory cortices. The nodal matching index (a measure of connection redundancy) was increased in the primary motor area and limb and trunk somatosensory areas. Increased matching index extended into retrosplenial and visual areas specifically in the onset phase. Interestingly, nodal eigenvector centrality was selectively reduced in the primary motor and somatosensory cortices throughout disease progression and increased in the retrosplenial cortex in the pre-disease and late phases.

These area-specific changes were expected as other disease models have shown area specific alterations in network configuration (21, 46, 50). What was unexpected was where the changes occurred, as we initially hypothesized that most network alterations would be concentrated in the sensorimotor regions as SCA8 in both patients and this mouse model has a significant motor component (5, 15). While many motor and sensory regions did show changes in network connectivity, major changes also occurred beyond the sensorimotor cortices in the visual and retrosplenial areas. The cerebellum has coherent activity with several neocortical regions including the barrel (51), prefrontal (52), primary motor, and somatosensory cortices (53, 54) which may explain changes outside the sensorimotor cortices. While these area-specific FC changes may result from the pathological and physiological changes in these mice, an alternative explanation is that the network changes are compensatory to preserve normal function. The increased connection strength and matching index may be a mechanism to maintain normal neural function in the sensorimotor system. While the mechanisms of cerebellar control of neocortical processing as well as SCA8’s impact on them remain unclear, our results demonstrate that profound alterations in neocortical processing occur in this canonically cerebellar disease.

### SCA8 genotype can be decoded using neocortical functional network alterations

Functional connectivity is being used as a biomarker for neurological diseases and diagnostic tool for disease progression (21, 24), including identifying early changes in presymptomatic individuals genetically positive for neurodegenerative diseases (55). Based on fMRI, FC has been used with machine learning to successfully classify individuals with major depressive disorder and show that alterations in the dorsal cingulate, prefrontal, and parietal cortices were the most influential areas for classification (56). Combined MRI imaging modalities can be used to distinguish pre-onset Huntington’s disease individuals from controls and predict years to disease onset (55). In childhood epilepsy, FC network metrics can be used to predict epilepsy duration using a machine learning algorithm (24).

Here, a stepwise GLM was used to predict animal genotype (SCA8+ or NT control) from the CCA matrices used to construct neural networks. The GLM decodes animal genotypes with >95% accuracy at all disease phases and when genotype data is shuffled, the predictive power of the GLM is lost. Furthermore, only a subset of connections is required for optimal decoding in each disease phase, most concentrated in the primary and accessory sensory cortices and higher visual integration areas. These data suggest that network integration is altered within and across sensory modalities that in turn contributes to motor dysfunction in SCA8 as mice are not able to efficiently process sensory information. Therefore, robust decoding results suggest network topology could be used to stage cognitive impairment and evaluate the efficacy of treatments in SCA8.

### Implications for network processing in SCA8

Any disruption of network homeostasis can lead to ill-configured information processing that correlates with disease state (24, 47). One explanation for SCA8+ FC shifting toward a globally hyperconnected state with increases in clustering of spatially distributed modules and efficiency is that the network has shifted to a more random topology where cross-regional integration has been diminished as hypothesized for HD and epilepsy (21, 24, 57). This explanation fits with the observed increases in global connection density and strength. This suggests an increase in the number of network shortcuts between distant regions which is also supported by the global increases in efficiency, centrality, transitivity, and number of communities. The result is a noisier network containing anatomically distributed communities resulting in degradation of information transfer and local processing specificity.

Our results suggesting an ill-configured network in SCA8 are consistent with disinhibition data obtained from computational models and bioengineered *in vitro* neural networks. In healthy macaque and cat cortical networks small-world topology, where spatially localized cliques of nodes/communities are densely connected within the grouping and more sparsely connected to other groups, dominates (58, 59). This network configuration allows efficient information transfer with minimal structural wiring costs and easier reprogramming by gating information transfer both within and between modules or communities. *In vitro* experiments demonstrate that modular networks are supportive of non-uniform conditional information transmission and dynamic transmission timing with inhibition providing powerful gating to transmission (60). Blockade of inhibition in *in vitro* networks (producing an aberrant state) dramatically increases information propagation and synchronization across network modules/communities causing the network to function uniformly as a single entity (60). The SCA8+ mice show properties similar to a disinhibited, more random network having global hyperconnectivity with increased connection strengths. As a result, it is likely that aberrant pathways of information transfer which are not functionally relevant are being potentiated leading to higher efficiency and an ill-configured anatomically distributed community structure.

### Toward a contributing mechanism to SCA8 pathogenesis

This study is the first showing long-range FC alterations in a mouse model of SCA8 at the mesoscale, a level between the microscopic and macroscopic. Both mouse models and human post-mortem tissue studies report cellular level alterations in SCA8 (2, 10, 15, 17, 61). These changes are likely due to RAN translation of pathological SCA8 RNA and sequestration of proteins such as MNBL1 (15), kelch-like protein 1 (KLHL1; 62), and other RNA binding proteins, in addition to abnormal protein aggregation in the cerebellum and other brain areas (2, 10, 17). Expression of pathogenic SCA8 alleles causes upregulation of GABT4, likely through reduced MBNL1 or increased CUGBP1 activity. As a result, the SCA8 cerebellum exhibits hyperexcitability due to increased reuptake of synaptic GABA and reduced inhibitory tone (2, 15). As many of the long-range connections between the cerebellar nuclei and other brain regions are glutamatergic, we propose that reduced inhibitory tone would increase excitatory cerebellar outputs and alter neocortical networks. Both our previous and current results support this hypothesis of global GABAergic dysfunction. A case study proposed a similar hypothesis after observing glucose hypometabolism in the cerebellum and neocortex, as well as reduced [11 C]-fluzamenil binding brain-wide in an SCA8 patient (63). These data suggest a globally altered GABAergic system contributes to SCA8 pathology and contributes to both cerebellar and extra-cerebellar symptoms of SCA8.

## Methods

### Sex as a biological variable

Our study examined both male and female mice. While a difference in age of onset was noted between sexes, changes in cortical networks for SCA8+ mice were similar, and sexes were combined for analyses.

### Animal model and surgical procedures

Transgenic mice (FVB) expressing the human SCA8 repeat expansion (SCA8+, 5 female, 2 male; Figure 1A, top) and non-transgenic FVB control animals (NTC; 5 female, 4 male) were used for this wide-field cortical Ca^2+^ imaging study (2). Briefly, mice 76 ± 23 (mean ± SD) days of age were anesthetized with isoflurane (5% induction; 1-2% maintenance) and implanted with transparent polymer skulls “See-Shells” and titanium headplates as previously described, including analgesia, monitoring, craniotomy, and post-operative care (28–31, 64). The craniotomy and polymer skull provided access to a large region of the dorsal neocortex. Images were taken of the craniotomy and cranial landmarks prior to skull removal. Immediately post-surgery, prior to recovery from anesthesia, animals were injected retro-orbitally with virally encoded GCaMP6f (AAV-PHP-eb-hSyn-GCaMP6f; 150 μL; titer: 3.81 x 10^12^ - 7 x 10^13^ gc/mL) which crosses the blood-brain barrier (65, 66). Post-surgery, mice were housed on a 12-hour reverse light-dark cycle and allowed 2 or more weeks (28 ± 12 days) recovery for sufficient viral expression and habituation to head-fixation on our treadmill. Post-experimental immunohistology for GCaMP6f expression in selected animals showed pan-neuronal infection of cells across all neocortical regions (Figure 1B, left) and neocortical cell layers (Figure 1B, right).

### Habituation and experimental setup

Mice were habituated to head-fixation and the disc treadmill over three sessions. In each session, mice were allowed to explore the treadmill freely for 5 minutes followed by increasing head-fixation time (5,10, and 20 minutes) over 3 days. If mice still showed signs of stress or discomfort during head-fixation, additional habituation sessions were given. As age of onset can vary dramatically in SCA8 (167 ± 47 days), imaging began around postnatal day 100 (103 ± 25 days) and mice were imaged weekly or biweekly up to 12 weeks post-onset or death (2). Imaging sessions took place during the animal’s dark phase and consisted of ∼10-12 trials (5.5 minutes long) per mouse per day. Imaging sessions were performed with mice head-fixed above the freely moving treadmill which allowed for awake resting, and a variety of motor behaviors (Figure 1A, bottom). Movement of the disc treadmill was monitored by a rotary encoder and recorded at 1 kHz by an Arduino microcontroller (Arduino Mega 2560; Arduino).

### Mesoscale Ca^2+^ imaging

Ca^2+^ imaging was performed with the animal head-fixed on the disc treadmill under a wide-field epifluorescence microscope (Nikon AZ-100; Nikon). Manual zoom was used to ensure the neocortex window maximally filled the imaging field (∼1.5 zoom). The microscope was focused below the brain surface to capture fluorescence from layers II/III of the neocortex (∼150-200 µm). Dual wavelength imaging was achieved by strobing LEDs using a switcher (OptoLED; Cairn) and alternating image frames of Ca^2+^-dependent GCaMP6f signal (470 nm) and Ca^2+^-independent signal (405 nm; Cairn) were captured using a high-speed CMOS camera (40 fps, 18 ms exposure; 256 x 256 pixels; Orca Flash4.0; Hamamatsu; ∼ 33.8 x 33.8 µm) and Metamorph software (Molecular Devices). Synchronization of the microscope camera as well as the rotary encoder was achieved using TTL pulses from a Power 1401 data acquisition system and Spike2 software (Cambridge Electronic Design).

### Processing of Ca^2+^ imaging data

Individual imaging trials for each mouse were pre-processed as previously described (28). Briefly, 470 and 405 nm images were deinterleaved and the first 30 seconds of each trial removed due to potential rundown of the Ca^2+^ signal. Ca^2+^-dependent GCaMP signals were corrected for Ca^2+^-independent GCaMP signals in a similar manner as prior studies (28, 31, 67, 68). For each experimental session, corrected Ca^2+^ images were saved along with a background (470 nm) image. A reference day was chosen for each mouse, a mask was drawn to separate out the neocortex field-of-view (FOV), and all imaging sessions were aligned to the reference day using standard Matlab registration functions (*imregconfig*; *imregtform*; *imwarp*). Registration of images within and between sessions was done to minimize artifacts due to slight changes in implant position in the imaging field and any motion (28).

As the number of imaging sessions per mouse was large and varied in number, we limited our analyses to three stages of SCA8 disease. The pre-disease phase was defined as the 2-3 imaging sessions (spanning 14 ± 2.2 days) prior to SCA8 onset. Disease onset was empirically defined as a greater than 10% reduction in maximum pre-onset body weight in SCA8+ mice (all: 167 ± 47 days; males: 156 ± 46 days; females: 184 ± 47 days; n = 15,6,9 respectively). For SCA8+ mice, having an NTC littermate in the same cohort, onset for the NTC littermate was at the same age as the SCA8+ littermate. For NTC animals without an SCA8+ littermate, onset was near the average onset age for the sex of the mouse (males: ∼150 days, females: ∼200 days). The onset phase of SCA8 was defined as the 2-3 imaging sessions (spanning 14 ± 1.8 days) after disease onset. The late disease phase of SCA8 was defined as the final 3 imaging sessions (spanning 13.9 ± 2.9 days) prior to death/euthanasia or the final 3 imaging sessions when the mouse reached 3 months post-onset (Figure 1C). For analyses, only imaging sessions within the 3 defined disease phases were used. The hemodynamic-corrected data for each mouse were spatiotemporally smoothed using a 3D 9x9x9 spatial gaussian filter (function: *imgaussfilt3*; Matlab 2022a) and concatenated. Signal noise due to LED illumination variability was removed by regressing the signal in the mask FOV against the signal in the neocortical FOV. Analysis was performed on the residuals of the regression. Denoised data were compressed using low rank singular value decomposition (SVD) keeping the first 200 components (28, 29, 32, 69).

### Spatial Independent Component Analysis

Spatial independent component analysis (ICA) was performed on the concatenated data set for each mouse, computing 60 independent components (ICs) using the Joint Approximation and Diagonalization of Eigenmatrices (JADE) algorithm that obtains maximally independent source signals from signal mixtures by minimizing mutual information (70, 71), as used in previous wide-field Ca^2+^ imaging studies (28, 29, 72). The solutions were multiplied back into the original vector space and z-scored to yield spatial maps of the ICs. Binary masks of the significant areas of the ICs were obtained by setting values between ± 2.5 SD to zero and all other values to one. Very small ICs with less than 250 contiguous pixels were excluded. Occasionally, ICs contained more than one region, for example a pair of homotopic regions. In these cases, the IC was separated, so each IC mask consisted of a single region. Remaining ICs were inspected for artifacts and were manually discarded, including areas overlying only vasculature (28, 29, 69).

To aid in results interpretation, the Allen Common Coordinate Framework (CCFv3) was aligned to the cranial window of each animal through a multi-step warping and alignment process using custom code modified from Paninski and colleagues (32, 33). For each animal, the surgical craniotomy image containing cranial landmarks (inferior cerebral vein near the frontonasal suture; bregma; midline; lambda) and full craniotomy drill path was cropped and rotated to match the orientation of the neocortical window during Ca^2+^ imaging sessions. The surgical image was used to align the craniotomy to the CCFv3 (32; Figure 1D) and the inverse transform was obtained to align the atlas to the final images. Next, the reference image for the Ca^2+^ imaging sessions was loaded. A mask was drawn around the drill path of the large craniotomy or the frame of the implant which sits directly above the drill path for each image, respectively (Figure 1E). The surgical craniotomy mask was registered to the implant frame mask (*imregconfig*; *imregtform*; *imwarp*) and the surgical craniotomy images containing cranial landmarks were used to align the craniotomy to the CCFv3 (32, 33). The resulting transformations were applied to the atlas aligning it to the reference image of the implant for each mouse. Once aligned, we were able to visualize ∼15 atlas regions in each hemisphere (30 total) of the mouse cortex (Figure 1F).

### Functional connectivity analysis

Using the aligned Allen CCF for each mouse, ICs were assigned to an atlas region based on position of their centers; or if on a border, which atlas region the IC overlapped with most. Functional connectivity (FC) adjacency matrices were computed across atlas regions using the average ΔF/F signal from ICs on a trial basis using canonical correlation (*canoncorr*), which gives a weighted correlation between sets of variables (in this case sets of ICs within two atlas regions). Only the first canonical correlation value was used in the FC matrices. To aid in comparison across genotypes, adjacency matrices for each mouse were expanded to include all possible atlas regions seen across mice (30 areas). When a particular atlas region was not represented for a mouse (did not have an IC assigned), zeros were placed in the adjacency matrix for that atlas region. To determine network structure and properties, expanded adjacency matrices were thresholded at canonical correlation values ≥ 0.5. FC graphs were plotted using the centroids of atlas regions as nodes and thresholded canonical correlation values as edges. The Brain Connectivity Toolbox was used to calculate FC graph properties at a trial level using thresholded FC matrices containing all 30 possible atlas areas, excluding strength, matching index, and community structure (34). Strength and matching index were calculated using unthresholded FC matrices. To calculate community structure (*Louvain-communities*), adjacency matrices containing only the represented nodes (atlas regions) for a mouse were used and the measure was subsequently normalized to the number of nodes in the graph.

To assess global network changes for measures calculated at the node level (such as strength and eigenvector centrality), values were averaged across nodes in each trial-based graph and subsequently averaged across animals. To calculate region-based changes in network properties, measures calculated at the node level (strength, centrality, degree, matching) were averaged for each node (i.e. all measures comparing the relevant node to other network nodes) and disease phase. While we were able to calculate some of the global measures at the node level (strength, eigenvector centrality), others are strictly global (density, global efficiency). To obtain similar measures at the node level, we determined nodal degree (as a proxy for connection density) and nodal matching index (to assess connectivity overlap between two nodes, with the assumption that more overlap suggests greater efficiency). Microsoft Excel (Microsoft Corporation, 2016), GraphPad Prism (Graphpad, 2024, Boston MA), and JMP Pro software (JMP Statistical Discovery LLC, 2024, Cary NC) were used to compare network properties across the three disease phases for SCA8+ mice and non-transgenic controls.

### GCaMP6f Immunohistochemistry and cortical expression

A selected set of retro-orbitally injected animals (both genotypes) were used for histology to verify the efficacy of viral expression. Mice were anesthetized with isoflurane (5%) and either perfused intracardially with 0.1 M phosphate buffered saline (PBS; pH ∼ 7.2-7.4) followed by paraformaldehyde (PFA; 4%) or injected intracardially with 0.3 mL Euthasol solution (Virbac). Brains were extracted and post-fixed for 1-3 days, then sectioned (50 μm sections; coronal) on either a cryostat or vibratome. For cryostat sectioning, brains were submerged in 30% sucrose solution 1-2 days prior to sectioning for dehydration. After sectioning, tissue was kept in antifreeze solution (30% glycerol; 30% ethylene glycol; 40% PBS) until needed for histology to prevent tissue degradation. At time of histology, tissue sections were washed with PBS (0.1 M; 3 times for 10 minutes) and blocked in a PBS solution containing 0.5% Triton-X 100 and 10% normal donkey serum (NDS; Sigma-Aldrich D9663) on an orbital shaker for 1 hour at room temperature. Tissue was incubated overnight at room temperature with a primary rabbit anti-GFP antibody (1:1000; A-6455 ThermoFisher). The following day, tissue was rinsed with PBS (0.1 M; 3 times for 10 minutes) and incubated with a donkey-anti-rabbit Alexa Fluor 555 secondary antibody (1:500; A-31572, ThermoFisher) for two hours at room temperature. Tissue was subsequently rinsed with PBS (0.1 M; 3 times for 10 minutes) and mounted with Invitrogen ProLong Diamond Antifade Mountant with DAPI (P36962, Invitrogen, ThermoFisher).

Tissue sections were imaged using a Leica Stellaris confocal microscope. Single plane tiled images of tissue sections were taken using a 20X objective lens (1024x1024 resolution) with the tunable excitation laser at 521 nm and an emission filter in the range of the Alexa Fluor 555 fluorescence peak (540-625 nm). Coronal neocortical images were collected, stitched, and merged using Leica LAS X software (Leica Microsystems). Cortical z-stacks were also acquired using a 20X objective with 2-3 frame averaging and an optical z-step of 1 µm. Images were processed using FiJi software (73). Tiled images or confocal stack maximum projections were background subtracted, denoised and contrast was enhanced for visualization.

### Statistical analysis

Data were analyzed and plotted using custom-written Matlab (*Matlab 2022a*) code and the Brain Connectivity Toolbox (34). For data presented in bar and line graph format as well as all tables, descriptive statistics shown are mean ± standard deviation. For data presented in violin plot format, descriptive statistics shown are median ± upper/lower quartile. Atlas maps showing average change in network measures per atlas area data are presented in a colorimetric manner with increases shown as red and decreases shown as blue (hue indicates change magnitude). Statistical analysis was performed using GraphPad Prism and JMP. Comparison of IC numbers and cortical coverage between genotypes were evaluated using non-parametric Mann-Whitney (MW) tests. Comparisons of global network measures across time and genotype were done at the trial level (∼450 trials per time point) using repeated measures mixed-effects two-way ANOVAs with post-hoc Bonferroni corrected pairwise comparisons.

Comparison of region-specific network measures was performed in JMP statistical software. A repeated measures mixed model 3-way ANOVA was done by fitting a random effects linear model to the data. Due to the size of the dataset at the trial level, measurements in finer atlas regions were pooled into major anatomical parcellations (primary motor cortex - MOp, secondary motor cortex - MOs, barrel fields - BFd, somatosensory cortices - SSp, retrosplenial cortex - RSP, visual and accessory visual cortices - VIS). The six major anatomical parcellations were used in the analysis of regional differences. However, we elected to show the magnitude of the changes in the FC measures to the subregions to provide a more granular spatial assessment of the region-specific differences. Post-hoc Tukey tests were used for pairwise comparisons between major regions and all subregions within each major region were considered significant regardless of the directionality of the differences.

## Study Approval

All experimental procedures were approved by the Institutional Care and Use Committee of the University of Minnesota.

## Data and Code Availability

The raw database of Ca^2+^ recordings in these animals consists of several terabytes of data. As such, the raw data will only be available upon request. Custom code will also be available upon request.

## Author Contributions

A.N., R.C., L.R., and T.E. conceptualized and designed the research. A.N, M.G., K.M conducted experiments. A.N., L.P., and R.C. analyzed the data. A.N., L.P., R.C., and T.E. interpreted the results of experiments and prepared figures. A.N., L.P., R.C., L.R., and T.E. provided supervision and guidance throughout the project. A.N., L.P., R.C., L.R., and T.E. wrote the manuscript.

## Acknowledgements

The authors and members of the Ebner lab would like to thank Lijuan Zhuo for assisting with rodent surgeries and general laboratory support during this project. We thank Evelyn Flaherty, William Chiesl, and Cecelia Huffman for their assistance in data collection. We thank the University of Minnesota’s Viral Vector Core, specifically Ezequiel Marron Fernandez de Velasco, for production of the viral vectors used in this study and the University of Minnesota’s Imaging Center for assistance with immunohistochemistry and tissue imaging. Finally, we thank all members of the Ebner and Ranum labs for their invaluable feedback during project execution and manuscript preparation. This work was funded in part by NIH grants P30 DA048742 (TJE), R01 NS111028 (TJE), RF1 NS126044 (TJE) and R37 NS040389 (LPWR).

## Notes

**Conflict of Interest Statement:** The authors have declared that there are no conflicts of interest.

### Competing Interest Statement

The authors have declared no competing interest.

## References

1. Castelli LM, Huang WP, Lin YH, Chang KY, and Hautbergue GM. Mechanisms of repeat-associated non-AUG translation in neurological microsatellite expansion disorders. Biochem Soc Trans. 2021;49(2):775–92.

2. Moseley ML, Zu T, Ikeda Y, Gao W, Mosemiller AK, Daughters RS, et al. Bidirectional expression of CUG and CAG expansion transcripts and intranuclear polyglutamine inclusions in spinocerebellar ataxia type 8. Nat Genet. 2006;38(7):758–69.

3. Moseley ML, Schut LJ, Bird TD, Koob MD, Day JW, and Ranum LP. SCA8 CTG repeat: en masse contractions in sperm and intergenerational sequence changes may play a role in reduced penetrance. Hum Mol Genet. 2000;9(14):2125–30.

4. Perez BA, Shorrock HK, Banez-Coronel M, Zu T, Romano LE, Laboissonniere LA, et al. CCG•CGG interruptions in high-penetrance SCA8 families increase RAN translation and protein toxicity. EMBO Mol Med. 2021;13(11):e14095.

5. Cleary JD, Subramony SH, and Ranum LPW. In: Adam MP, Feldman J, Mirzaa GM, Pagon RA, Wallace SE, Bean LJH, et al. eds. GeneReviews(®). Seattle (WA): University of Washington, Seattle Copyright © 1993-2024, University of Washington, Seattle. GeneReviews is a registered trademark of the University of Washington, Seattle. All rights reserved.; 1993.

6. Day JW, Schut LJ, Moseley ML, Durand AC, and Ranum LP. Spinocerebellar ataxia type 8: clinical features in a large family. Neurology. 2000;55(5):649–57.

7. Lilja A, Hämäläinen P, Kaitaranta E, and Rinne R. Cognitive impairment in spinocerebellar ataxia type 8. J Neurol Sci. 2005;237(1-2):31–8.

8. Ikeda Y, Shizuka M, Watanabe M, Okamoto K, and Shoji M. Molecular and clinical analyses of spinocerebellar ataxia type 8 in Japan. Neurology. 2000;54(4):950–5.

9. Juvonen V, Hietala M, Päivärinta M, Rantamäki M, Hakamies L, Kaakkola S, et al. Clinical and genetic findings in Finnish ataxia patients with the spinocerebellar ataxia 8 repeat expansion. Ann Neurol. 2000;48(3):354–61.

10. Zu T, Gibbens B, Doty NS, Gomes-Pereira M, Huguet A, Stone MD, et al. Non-ATG-initiated translation directed by microsatellite expansions. Proc Natl Acad Sci U S A. 2011;108(1):260–5.

11. Zu T, Pattamatta A, and Ranum LPW. Repeat-Associated Non-ATG Translation in Neurological Diseases. Cold Spring Harb Perspect Biol. 2018;10(12).

12. Cleary JD, and Ranum LP. Repeat-associated non-ATG (RAN) translation in neurological disease. Hum Mol Genet. 2013;22(R1):R45–51.

13. Goodwin M, Mohan A, Batra R, Lee KY, Charizanis K, Fernández Gómez FJ, et al. MBNL Sequestration by Toxic RNAs and RNA Misprocessing in the Myotonic Dystrophy Brain. Cell Rep. 2015;12(7):1159–68.

14. Sta Maria NS, Zhou C, Lee SJ, Valiulahi P, Li X, Choi J, et al. Mbnl1 and Mbnl2 regulate brain structural integrity in mice. Commun Biol. 2021;4(1):1342.

15. Daughters RS, Tuttle DL, Gao W, Ikeda Y, Moseley ML, Ebner TJ, et al. RNA gain-of-function in spinocerebellar ataxia type 8. PLoS Genet. 2009;5(8):e1000600.

16. Zeman A, Stone J, Porteous M, Burns E, Barron L, and Warner J. Spinocerebellar ataxia type 8 in Scotland: genetic and clinical features in seven unrelated cases and a review of published reports. J Neurol Neurosurg Psychiatry. 2004;75(3):459–65.

17. Ayhan F, Perez BA, Shorrock HK, Zu T, Banez-Coronel M, Reid T, et al. SCA8 RAN polySer protein preferentially accumulates in white matter regions and is regulated by eIF3F. EMBO J. 2018;37(19).

18. Bürk K, Globas C, Bösch S, Klockgether T, Zühlke C, Daum I, et al. Cognitive deficits in spinocerebellar ataxia type 1, 2, and 3. J Neurol. 2003;250(2):207–11.

19. Duarte JV, Faustino R, Lobo M, Cunha G, Nunes C, Ferreira C, et al. Parametric fMRI of paced motor responses uncovers novel whole-brain imaging biomarkers in spinocerebellar ataxia type 3. Hum Brain Mapp. 2016;37(10):3656–68.

20. Bares M, Lungu OV, Liu T, Waechter T, Gomez CM, and Ashe J. The neural substrate of predictive motor timing in spinocerebellar ataxia. Cerebellum. 2011;10(2):233–44.

21. Harrington DL, Rubinov M, Durgerian S, Mourany L, Reece C, Koenig K, et al. Network topology and functional connectivity disturbances precede the onset of Huntington’s disease. Brain. 2015;138(8):2332–46.

22. Vasilkovska T, Adhikari MH, Van Audekerke J, Salajeghe S, Pustina D, Cachope R, et al. Resting-state fMRI reveals longitudinal alterations in brain network connectivity in the zQ175DN mouse model of Huntington’s disease. Neurobiol Dis. 2023;181:106095.

23. Bernhardt BC, Bonilha L, and Gross DW. Network analysis for a network disorder: The emerging role of graph theory in the study of epilepsy. Epilepsy Behav. 2015;50:162–70.

24. Paldino MJ, Zhang W, Chu ZD, and Golriz F. Metrics of brain network architecture capture the impact of disease in children with epilepsy. Neuroimage Clin. 2017;13:201–8.

25. Henschke JU, and Pakan JM. Disynaptic cerebrocerebellar pathways originating from multiple functionally distinct cortical areas. Elife. 2020;9.

26. Suzuki L, Coulon P, Sabel-Goedknegt EH, and Ruigrok TJ. Organization of cerebral projections to identified cerebellar zones in the posterior cerebellum of the rat. J Neurosci. 2012;32(32):10854–69.

27. Fujita H, Kodama T, and du Lac S. Modular output circuits of the fastigial nucleus for diverse motor and nonmotor functions of the cerebellar vermis. Elife. 2020;9.

28. Nietz AK, Streng ML, Popa LS, Carter RE, Flaherty EB, Aronson JD, et al. To be and not to be: wide-field Ca2+ imaging reveals neocortical functional segmentation combines stability and flexibility. Cereb Cortex. 2023.

29. West SL, Aronson JD, Popa LS, Feller KD, Carter RE, Chiesl WM, et al. Wide-Field Calcium Imaging of Dynamic Cortical Networks during Locomotion. Cereb Cortex. 2022;32(12):2668–87.

30. West SL, Gerhart ML, and Ebner TJ. Wide-field calcium imaging of cortical activation and functional connectivity in externally- and internally-driven locomotion. bioRxiv. 2023:2023.04.10.536261.

31. Cramer SW, Haley SP, Popa LS, Carter RE, Scott E, Flaherty EB, et al. Wide-field calcium imaging reveals widespread changes in cortical functional connectivity following mild traumatic brain injury in the mouse. Neurobiol Dis. 2023;176:105943.

32. Saxena S, Kinsella I, Musall S, Kim SH, Meszaros J, Thibodeaux DN, et al. Localized semi-nonnegative matrix factorization (LocaNMF) of widefield calcium imaging data. PLoS Comput Biol. 2020;16(4):e1007791.

33. Wang Q, Ding S-L, Li Y, Royall J, Feng D, Lesnar P, et al. The Allen mouse brain common coordinate framework: a 3D reference atlas. Cell. 2020;181(4):936–53. e20.

34. Rubinov M, and Sporns O. Complex network measures of brain connectivity: uses and interpretations. Neuroimage. 2010;52(3):1059–69.

35. Fornito A, Zalesky A, and Bullmore E. Fundamentals of brain network analysis. Academic press; 2016.

36. Koutrouli M, Karatzas E, Paez-Espino D, and Pavlopoulos GA. A Guide to Conquer the Biological Network Era Using Graph Theory. Front Bioeng Biotechnol. 2020;8:34.

37. White BR, Bauer AQ, Snyder AZ, Schlaggar BL, Lee J-M, and Culver JP. Imaging of functional connectivity in the mouse brain. PloS one. 2011;6(1):e16322.

38. Cramer JV, Gesierich B, Roth S, Dichgans M, Düring M, and Liesz A. In vivo widefield calcium imaging of the mouse cortex for analysis of network connectivity in health and brain disease. Neuroimage. 2019;199:570–84.

39. Bero AW, Bauer AQ, Stewart FR, White BR, Cirrito JR, Raichle ME, et al. Bidirectional relationship between functional connectivity and amyloid-β deposition in mouse brain. J Neurosci. 2012;32(13):4334–40.

40. Kumar N, and Miller GM. White matter hyperintense lesions in genetically proven spinocerebellar ataxia 8. Clinical Neurology and Neurosurgery. 2008;110(1):65–8.

41. Bezzina C, Verret L, Juan C, Remaud J, Halley H, Rampon C, et al. Early onset of hypersynchronous network activity and expression of a marker of chronic seizures in the Tg2576 mouse model of Alzheimer’s disease. PloS one. 2015;10(3):e0119910.

42. Kazim SF, Chuang S-C, Zhao W, Wong RKS, Bianchi R, and Iqbal K. Early-Onset Network Hyperexcitability in Presymptomatic Alzheimer’s Disease Transgenic Mice Is Suppressed by Passive Immunization with Anti-Human APP/Aβ Antibody and by mGluR5 Blockade. Frontiers in Aging Neuroscience. 2017;9.

43. Cocozza S, Costabile T, Tedeschi E, Abate F, Russo C, Liguori A, et al. Cognitive and functional connectivity alterations in Friedreich’s ataxia. Annals of Clinical and Translational Neurology. 2018;5(6):677–86.

44. Guo J, Jiang Z, Liu X, Li H, Biswal BB, Zhou B, et al. Cerebello-cerebral resting-state functional connectivity in spinocerebellar ataxia type 3. Human Brain Mapping. 2023;44(3):927–36.

45. Blaschke SJ, Hensel L, Minassian A, Vlachakis S, Tscherpel C, Vay SU, et al. Translating Functional Connectivity After Stroke: Functional Magnetic Resonance Imaging Detects Comparable Network Changes in Mice and Humans. Stroke. 2021;52(9):2948–60.

46. Lorenzini L, Ingala S, Collij LE, Wottschel V, Haller S, Blennow K, et al. Eigenvector centrality dynamics are related to Alzheimer’s disease pathological changes in non-demented individuals. Brain Communications. 2023;5(3).

47. Werner CJ, Dogan I, Saß C, Mirzazade S, Schiefer J, Shah NJ, et al. Altered resting-state connectivity in Huntington’s disease. Human brain mapping. 2014;35(6):2582–93.

48. Burgstaller J, Hindinger E, Donovan J, Dal Maschio M, Kist AM, Gesierich B, et al. Light-sheet imaging and graph analysis of antidepressant action in the larval zebrafish brain network. bioRxiv. 2019:618843.

49. Nelson CJ, and Bonner S. Neuronal Graphs: A Graph Theory Primer for Microscopic, Functional Networks of Neurons Recorded by Calcium Imaging. Frontiers in Neural Circuits. 2021;15.

50. Wang J, Zuo X, Dai Z, Xia M, Zhao Z, Zhao X, et al. Disrupted Functional Brain Connectome in Individuals at Risk for Alzheimer’s Disease. Biological Psychiatry. 2013;73(5):472–81.

51. O’Connor SM, Berg RW, and Kleinfeld D. Coherent Electrical Activity Between Vibrissa Sensory Areas of Cerebellum and Neocortex Is Enhanced During Free Whisking. Journal of Neurophysiology. 2002;87(4):2137–48.

52. Watson TC, Becker N, Apps R, and Jones MW. Back to front: cerebellar connections and interactions with the prefrontal cortex. Front Syst Neurosci. 2014;8:4.

53. Proville RD, Spolidoro M, Guyon N, Dugué GP, Selimi F, Isope P, et al. Cerebellum involvement in cortical sensorimotor circuits for the control of voluntary movements. Nat Neurosci. 2014;17(9):1233–9.

54. Heck DH, Fox MB, Correia Chapman B, McAfee SS, and Liu Y. Cerebellar control of thalamocortical circuits for cognitive function: A review of pathways and a proposed mechanism. Frontiers in Systems Neuroscience. 2023;17.

55. Rizk-Jackson A, Stoffers D, Sheldon S, Kuperman J, Dale A, Goldstein J, et al. Evaluating imaging biomarkers for neurodegeneration in pre-symptomatic Huntington’s disease using machine learning techniques. Neuroimage. 2011;56(2):788–96.

56. Geng X, Xu J, Liu B, and Shi Y. Multivariate classification of major depressive disorder using the effective connectivity and functional connectivity. Frontiers in neuroscience. 2018;12:38.

57. Van Den Heuvel MP, and Sporns O. Rich-club organization of the human connectome. Journal of Neuroscience. 2011;31(44):15775–86.

58. Sporns O, Tononi G, and Edelman GM. Connectivity and complexity: the relationship between neuroanatomy and brain dynamics. Neural Networks. 2000;13(8):909–22.

59. Achard S, and Bullmore E. Efficiency and cost of economical brain functional networks. PLoS computational biology. 2007;3(2):e17.

60. Shein-Idelson M, Cohen G, Ben-Jacob E, and Hanein Y. Modularity induced gating and delays in neuronal networks. PLoS Computational Biology. 2016;12(4):e1004883.

61. He Y, Zu T, Benzow KA, Orr HT, Clark HB, and Koob MD. Targeted deletion of a single Sca8 ataxia locus allele in mice causes abnormal gait, progressive loss of motor coordination, and Purkinje cell dendritic deficits. Journal of Neuroscience. 2006;26(39):9975–82.

62. Nemes JP, Benzow KA, and Koob MD. The SCA8 transcript is an antisense RNA to a brain-specific transcript encoding a novel actin-binding protein (KLHL1). Human molecular genetics. 2000;9(10):1543–51.

63. Terada T, Kono S, Konishi T, Miyajima H, and Ouchi Y. Altered GABAergic system in the living brain of a patient with spinocerebellar ataxia type 8. Journal of neurology. 2013;260:3164–6.

64. Ghanbari L, Carter RE, Rynes ML, Dominguez J, Chen G, Naik A, et al. Cortex-wide neural interfacing via transparent polymer skulls. Nature communications. 2019;10(1):1500.

65. Yardeni T, Eckhaus M, Morris HD, Huizing M, and Hoogstraten-Miller S. Retro-orbital injections in mice. Lab animal. 2011;40(5):155–60.

66. Chan KY, Jang MJ, Yoo BB, Greenbaum A, Ravi N, Wu W-L, et al. Engineered AAVs for efficient noninvasive gene delivery to the central and peripheral nervous systems. Nature neuroscience. 2017;20(8):1172–9.

67. Jacobs EA, Steinmetz NA, Peters AJ, Carandini M, and Harris KD. Cortical state fluctuations during sensory decision making. Current Biology. 2020;30(24):4944–55. e7.

68. MacDowell CJ, and Buschman TJ. Low-dimensional spatiotemporal dynamics underlie cortex-wide neural activity. Current Biology. 2020;30(14):2665–80. e8.

69. Musall S, Kaufman MT, Juavinett AL, Gluf S, and Churchland AK. Single-trial neural dynamics are dominated by richly varied movements. Nat Neurosci. 2019;22(10):1677–86.

70. Cardoso J-F. High-order contrasts for independent component analysis. Neural computation. 1999;11(1):157–92.

71. Sahonero-Alvarez G, and Calderon H. Proceedings of the 8th International Multi-Conference on Complexity, Informatics and Cybernetics. 2017:17–22.

72. Makino H, Ren C, Liu H, Kim AN, Kondapaneni N, Liu X, et al. Transformation of cortex-wide emergent properties during motor learning. Neuron. 2017;94(4):880–90 e8.

73. Schindelin J, Arganda-Carreras I, Frise E, Kaynig V, Longair M, Pietzsch T, et al. Fiji: an open-source platform for biological-image analysis. Nat Methods. 2012;9(7):676–82.

